# Persistent contacts between Climp63 and microtubules cause mitotic defects and nuclear fragmentation

**DOI:** 10.1101/2025.09.01.673496

**Authors:** Jelmi uit de Bos, Ulrike Kutay

## Abstract

The endoplasmic reticulum (ER) is the most elaborate endomembrane system in mammalian cells. To generate its characteristic shape, ER-resident proteins directly affect membrane topology, while interactions with the microtubule (MT) network determine ER positioning. The ER undergoes continuous remodeling with the most drastic rearrangements taking place during mitosis. A critical aspect is the reconfiguration of MT-ER interactions that accompanies spindle formation. One abundant ER-MT tether is constituted by the ER sheet-associated membrane protein Climp63, which binds MTs in a phosphorylation-sensitive manner. Here, we demonstrate that expression of a phosphodeficient Climp63 mutant in HeLa cells prevents dissolving of ER-MT contacts in mitosis, resulting in severe mitotic delays. Moreover, during mitotic exit, cells fail to properly enclose all chromosomes into a single nucleus, leading to excessive nuclear fragmentation. The emerging micronuclei assemble NE components with nuclear pore complexes, in contrast to what is known for micronuclei formed from lagging chromosomes. We further show that the N-terminal 28 amino acids are sufficient for the interaction between Climp63 and MTs with phosphorylation of serine S17 by CDK1 being critical for the mitotic release of the ER from MTs. Overall, our results demonstrate that aberrant Climp63 activity severely affects mitosis. One may speculate that mitotic aberrations may contribute to the poor prognosis associated with Climp63 overexpressing cancers.

## Introduction

The endoplasmic reticulum (ER) is a highly interconnected membrane network that is composed of functionally and morphologically distinct domains, including the nuclear envelope (NE) and the peripheral ER (Obara et al., 2023). Its wide range of morphologies are dependent on a set of ER-intrinsic shaping proteins that can induce high or low membrane curvatures or induce membrane fusion, thereby generating a continuous membrane network comprised of sheets, tubules and intermediate morphologies (Goyal & Blackstone, 2013, Shibata et al., 2006, Westrate et al., 2015). In addition, the microtubule (MT) network plays a pivotal role in determining ER network topology and distribution, which affects both ER tubules and sheets. On the one hand, almost all ER tubules are formed with aid of the MT network (Guo et al., 2018). On the other hand, ER sheet-shaping proteins interact with specific populations of MTs to regulate the global distribution of the ER throughout the cytoplasm (Zheng et al., 2022).

When metazoan cells prepare for open mitosis, changes in ER-MT interactions facilitate drastic membrane remodeling that affects both the NE and the peripheral ER (Deolal et al., 2024, Kors & Schlaitz, 2024). Although the mitotic ER remains a continuous network (Ellenberg et al., 1997, Yang et al., 1997), the NE as a distinctive ER subdomain is dissolved with the help of dynein-dependent MT forces (Beaudouin et al., 2002, Salina et al., 2002, Turgay et al., 2014). In addition, reduced contacts between the ER and the MT network lead to further morphological alterations of the mitotic ER (Deolal et al., 2024, Kors & Schlaitz, 2024). Several studies have investigated the consequences of the failed mitotic remodeling of specific MT-ER tubule interfaces (Chung et al., 2016, Nourbakhsh et al., 2021, Schlaitz et al., 2013, Smyth et al., 2012), highlighting the importance of remodeling ER tubule-MT interactions for mitotic fidelity. However, it is currently unknown how failures in remodeling the ER sheet-MT interface affects mitotic progression and chromosome segregation.

Climp63, also known as CKAP4, is a highly abundant (Itzhak et al., 2016, Uhlen et al., 2015), non-glycosylated type II transmembrane protein that is a key member of the ER sheet-MT interface. It is best known for its role in ER sheet formation, with its overexpression leading to an increase in the relative number of ER sheets (Parlakgul et al., 2022, Sandoz & van der Goot, 2015, Shibata et al., 2010, Wang et al., 2022, Zhao et al., 2023). Previous biochemical analyses have suggested that the majority of Climp63 is in a trimeric state (Sandoz et al., 2023). AlphaFold predicts that its luminal domain forms an extended trimeric coiled-coil, whereas the cytoplasmic domain is largely unstructured (Figure 1A). The cytoplasmic domain of Climp63 is required for interactions with the MT network and is phosphorylated in mitosis, but how Climp63 binds to MTs is debated (Klopfenstein et al., 1998, Vedrenne et al., 2005, Zheng et al., 2022).

**Figure 1.**
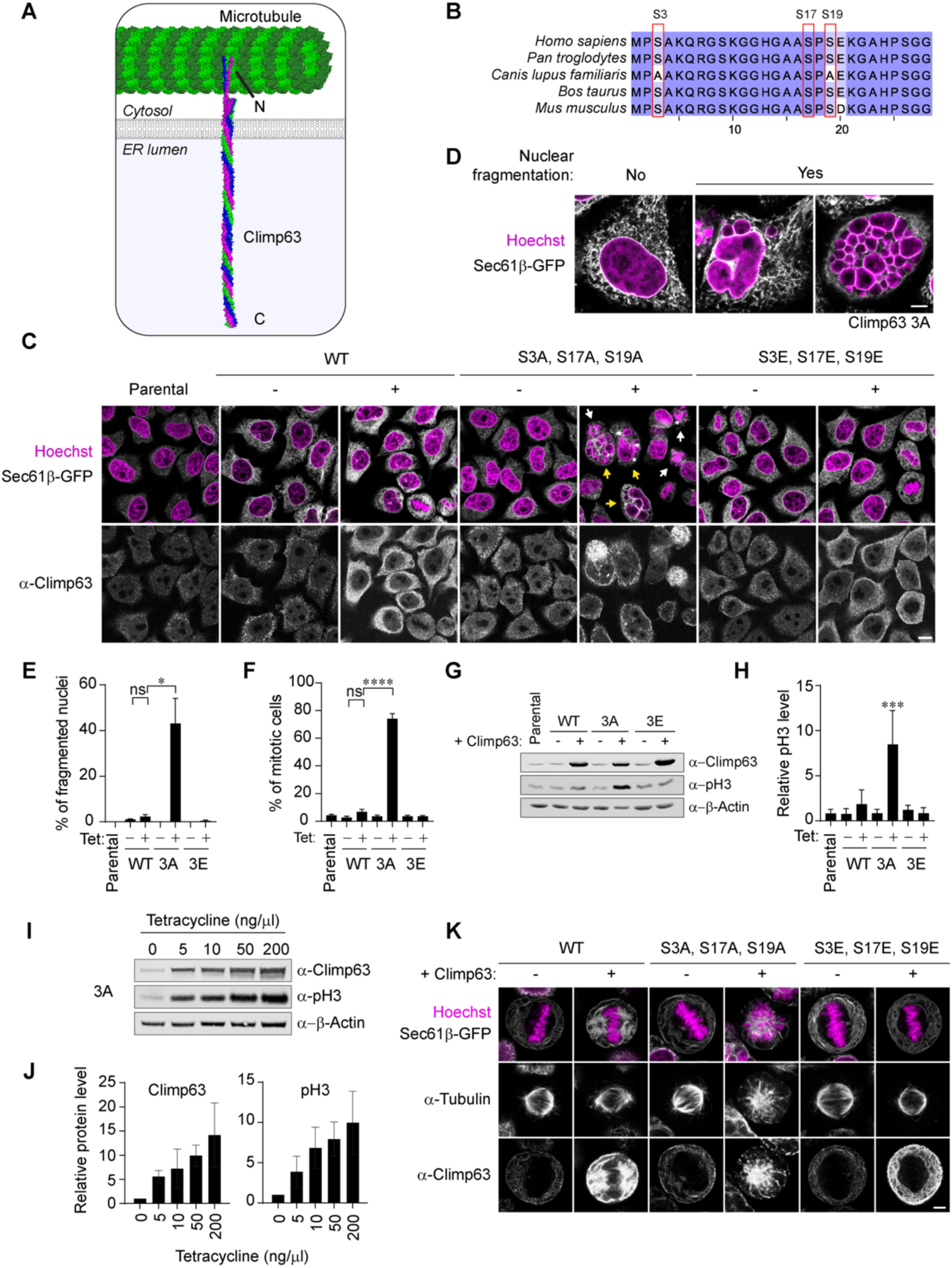
Expression of a phosphodeficient Climp63 mutant results in nuclear fragmentation and prolonged mitosis. **(A)** Model of full-length trimeric Climp63 in the ER membrane and its interaction with a microtubule (structure based on PDB 7SJ9). The structure of Climp63 was adapted from an AlphaFold prediction. **(B)** Multi-sequence alignment of the first 28 amino acids of Climp63 for the indicated species generated by JalView2 (Waterhouse et al., 2009). Experimentally confirmed phosphorylation sites are highlighted by red boxes. Color intensity reflects the degree of conservation. **(C)** Immunofluorescence analysis of tetracycline-inducible HeLa cell lines expressing Climp63 WT, 3A (S3A, S17A, S19A), or 3E (S3E, S17E, S19E) for 48 h. Sec61β-GFP as an ER marker is stably integrated into these cell lines. White arrows point out mitotic cells that are frequently observed upon 3A expression, yellow arrows point out cells with nuclear fragmentation. Scale bar, 10 µm, N = 3. **(D)** Representative images of nuclear fragmentation induced by the expression of phosphodeficient Climp63 in HeLa cells and their classification from the experiment in (C). Scale bar, 5 µm. **(E)** Quantification of the fraction of interphase cells with nuclear fragmentation. N = 3, n ≥ 90, mean ± SEM, *p ≤ 0.05, ns = not significant. **(F)** Quantification of the percentage of mitotic cells. N = 3, n ≥ 425, mean ± SEM, ****p ≤ 0.0001, ns = not significant. **(G)** Immunoblot analysis of samples from (C) with the indicated antibodies. N = 3. **(H)** Quantification of phospho-H3 (pH3) levels in (G) normalized to β-Actin and to the mean within replicates. Asterisks denote significant differences compared to the parental control. Mean ± SD, ***p ≤ 0.001. **(I)** Immunoblot analysis of phospho-H3 (pH3) levels after Climp63 3A induction with different tetracycline concentrations for 48 h. N = 3. **(J)** Quantification of protein levels normalized to β-Actin and the uninduced condition from three replicates of (H). Mean ± SD. **(K)** Representative images of immunofluorescence analysis of HeLa cells in metaphase expressing Climp63 WT, 3A or 3E and stained with the indicated antibodies. Scale bar, 5 µm.

Since Climp63 is an abundant tether of ER sheets to the MT network, we set out to explore how Climp63 is dissociated from MTs during mitotic onset and how defects in dissociation affect mitotic progression and fidelity. Here, we show that the expression of a phosphodeficient Climp63 mutant leads to massive mitotic defects manifesting in severe nuclear fragmentation in postmitotic cells. Live cell microscopy revealed that persistent, Climp63-mediated mitotic MT-ER interactions lead to mislocalization of the ER to the mitotic spindle. ER attachment to the mitotic spindle interferes with chromosome alignment at the metaphase plate, leading to a strong mitotic delay by activation of the spindle assembly checkpoint. Using the mitotic defects induced by expression of phosphodeficient Climp63 as a readout for MT binding, we demonstrate that the N-terminal region comprising aa 1-28 is sufficient for MT binding of Climp63 *in vivo* and pinpoint a single CDK1 phosphorylation site, S17, to be decisive for breaking MT-ER interactions during mitotic entry. Collectively, our results highlight the importance of dissolving ER-MT interactions for mitotic fidelity and shed new insights onto the role of the ER in mitosis and how persistent ER-spindle interactions affect mitotic progression.

## Results

### Expression of a phosphodeficient Climp63 mutant leads to mitotic defects

To investigate the role of Climp63-dependent MT-ER interactions in mitosis, we investigated three phosphorylation sites that are thought to regulate MT binding (S3, S17 and S19; Figure 1B) (Vedrenne et al., 2005). To this end, we generated HeLa cell lines that allow the tetracycline-inducible expression of either wild-type (WT), phosphodeficient (S3A, S17A, S19A) or phosphomimetic (S3E, S17E, S19E) Climp63 variants. Immunofluorescence analysis showed a striking increase in the number of both mitotic cells and interphase cells with fragmented nuclei 48 h after induction of the phosphodeficient Climp63 (3A) mutant (Figure 1C, D). The observed nuclear fragmentation is severe, frequently giving rise to more than 30 micronuclei. In contrast, cells expressing either WT or phosphomimetic Climp63 (3E) did not display mitotic errors. Quantification showed that over 75% of cells expressing phosphodeficient Climp63 are in mitosis. Of the remaining 25% of cells, more than 40% show some degree of nuclear fragmentation (Figure 1E, F). We also immunoblotted for phosphorylated histone H3 (S10), a mitosis-specific histone mark (Fischle et al., 2005, Wilkins et al., 2014). As expected, its levels are increased upon expression of Climp63 3A compared to the WT or 3E control (Figure 1G, H).

To exclude that the mitotic phenotype caused by Climp63 3A expression is due to higher expression levels of the phosphodeficient mutant as compared to the wildtype and phosphomimetic variants, we checked Climp63 levels and confirmed that Climp63 levels in all three cell lines were similar upon tetracycline induction, demonstrating that this phenotype is related to the mutations and not expression levels. Of course, expression levels of Climp63 did exceed those of the parental and uninduced cell lines. We therefore performed a tetracycline-titration series of the Climp63 3A cell line to investigate how expression levels affect the number of mitotic cells. Immunoblotting showed that increasing levels of Climp63 3A lead to a progressively stronger mitotic delay (Figure 1I, J). Together, these results indicate a failure in mitosis upon expression of phosphodeficient Climp63 that leads to severe nuclear fragmentation.

Given the mitotic defects in the presence of Climp63 3A, we set out to examine mitotic cells more carefully by immunofluorescence analysis. Interestingly, this revealed that the mitotic spindle of cells expressing Climp63 3A was distorted and aberrantly positioned (Figure 1K). We also failed to observe a clear bipolar orientation of spindles. In addition, the ER marker Sec61β-GFP revealed that the ER was repositioned from its normal peripheral localization and instead colocalized with the mitotic spindle. Together, this suggests that regulating Climp63 activity by phosphorylation is needed to spatially separate the ER from the mitotic spindle and that this is key for mitotic progression.

### Climp63 S17 regulates MT binding and is phosphorylated by CDK1

Given the persistent interaction of the ER with MTs upon mutation of the three phosphorylation sites in Climp63, we wished to understand which of these sites contribute to the regulation of MT binding in mitosis. To this end, we generated tetracycline-inducible phosphodeficient mutants at individual sites (S3A, S17A and S19A). We then performed immunofluorescence analysis of these cell lines and compared them to the WT and the 3A mutant as negative and positive controls, respectively. Akin to the Climp63 3A mutant, we observed a clear increase in the number of mitotic cells and of cells with nuclear fragmentation upon expression of Climp63 S17A but not for the S3A or S19A mutants (Figure 2A, C, D). Immunoblotting showed that Climp63 levels were similar upon tetracycline induction in all cell lines, while the proportion of mitotic cells specifically increased for the Climp63 S17A single mutant as illustrated by the phospho-H3 signal (Figure 2B). Together, these data point at a critical role for site S17 in the regulation of MT binding in Climp63 during mitosis.

**Figure 2.**
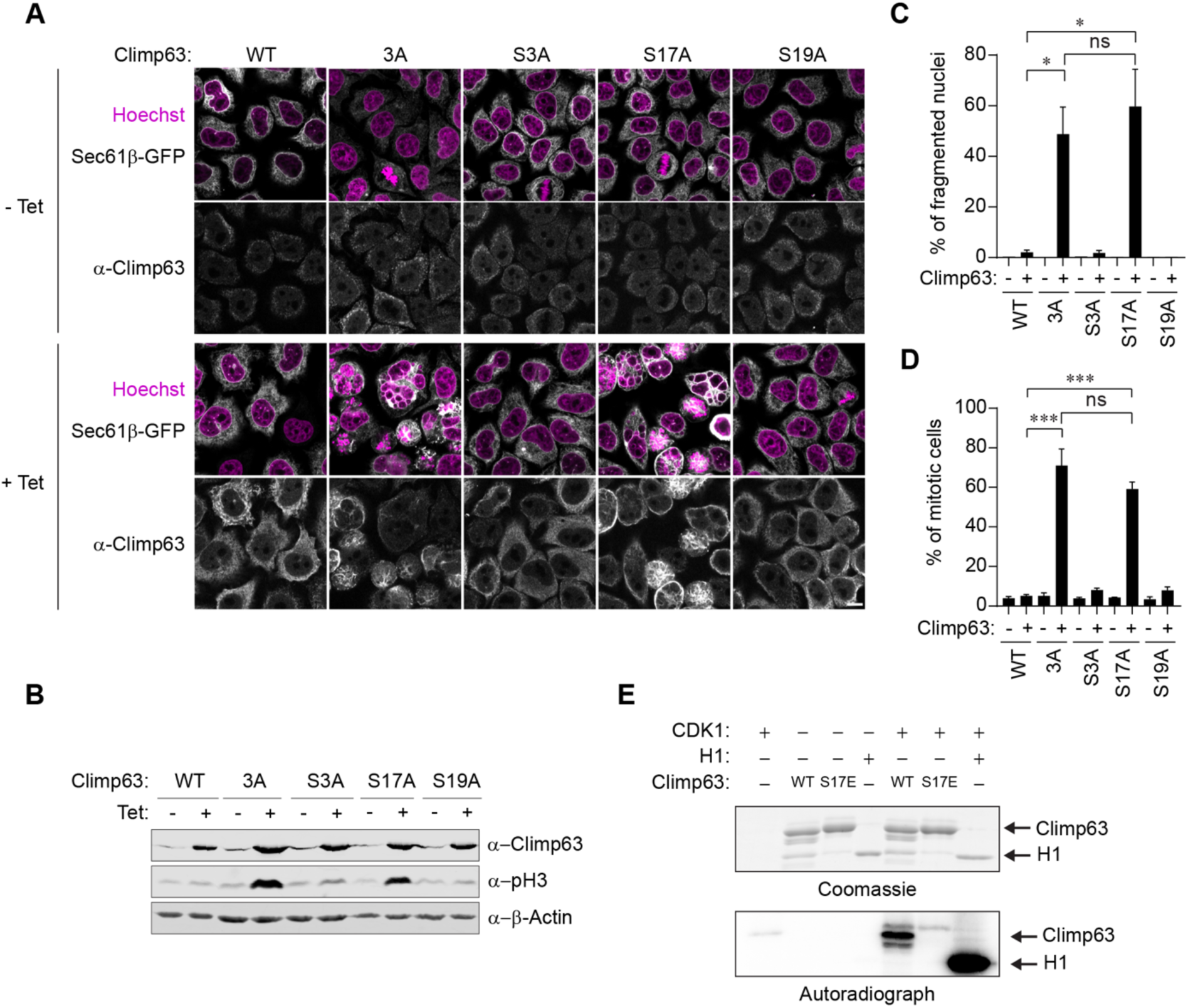
Phosphosite S17A is important to release ER from microtubules in mitosis and is a target of CDK1. **(A)** Immunofluorescence analysis of HeLa cell lines expressing Climp63 mutants that are rendered phosphorylation-deficient at individual phosphosites compared to the WT and 3A mutant. Sec61β-GFP as an ER marker is stably integrated into these cell lines. Scale bar, 10 µm. N = 3. **(B)** Immunoblot analysis of the same samples as in (A) with the indicated antibodies. N = 3. **(C)** Quantification of the percentage of interphase cells with nuclear fragmentation. N = 3, n ≥ 120, mean ± SEM, *p ≤ 0.05. **(D)** Quantification of the percentage of mitotic cells. N = 3, n ≥ 420, mean ± SEM. ***p ≤ 0.001. **(E)** *In vitro* kinase assay with recombinant purified Cyclin B1-CDK1 and Climp63 1-31-GFP-HA WT or S17E in the presence of γ-[32P]ATP. Histone H1 phosphorylation was used as positive control. N = 3.

Climp63 S17 is a proline-directed phosphorylation site, which is a common feature of CDK1 substrates. To test if Climp63 S17 could be a substrate of CDK1, we purified either a WT or a phosphomimetic (S17E) Climp63 fragment (1-31) and performed an *in vitro* phosphorylation assay using recombinant Cyclin B1-CDK1. Incubation of the kinase complex with the positive control histone H1 and the WT fragment showed incorporation of ^32^P phosphate, whereas the S17E mutant was not phosphorylated (Figure 2E). This confirmed that CDK1 can indeed phosphorylate Climp63 at site S17 *in vitro*.

### The N-terminal 28 residues of Climp63 are sufficient for MT-ER interactions

Currently, no structural data is available for the direct interaction of ER-shaping proteins and MTs. Indirect interactions have been characterized and include binding of SxIP motifs present in STIM1 and TAOK2 to the MT +TIP associated protein EB1 (Grigoriev et al., 2008, Honnappa et al., 2009, Nourbakhsh et al., 2021). Climp63 does not contain any SxIP motifs and is assumed to interact with MTs directly (Klopfenstein et al., 1998). The MT binding region of Climp63 has been studied before, but these studies have yielded inconclusive and partially contradictory results (Klopfenstein et al., 1998, Vedrenne et al., 2005, Zheng et al., 2022). With the aim to study the MT binding regions of the cytoplasmic domain of Climp63, we expressed the cytoplasmic domain of Climp63 (1-106) with a C-terminal HA tag in HeLa cells and performed immunofluorescence analysis (Figure 3A). We failed to observe any colocalization between Climp63 1-106 and MTs, suggesting that the affinity of the cytoplasmic domain by itself is not high enough to bind to MTs. We note that this observation is in contrast to another study that reported a direct interaction of Climp63 with MTs and showed complete colocalization of Climp63 1-106 with MTs (Klopfenstein et al., 1998), but our conclusion is supported by a later publication that reported results similar to ours (Nikonov et al., 2007). This would suggest that either the membrane association of Climp63 provides the avidity needed to bind MTs or that the interaction is not direct as suggested before (Klopfenstein et al., 1998), and instead facilitated by another (membrane-bound) component.

**Figure 3.**
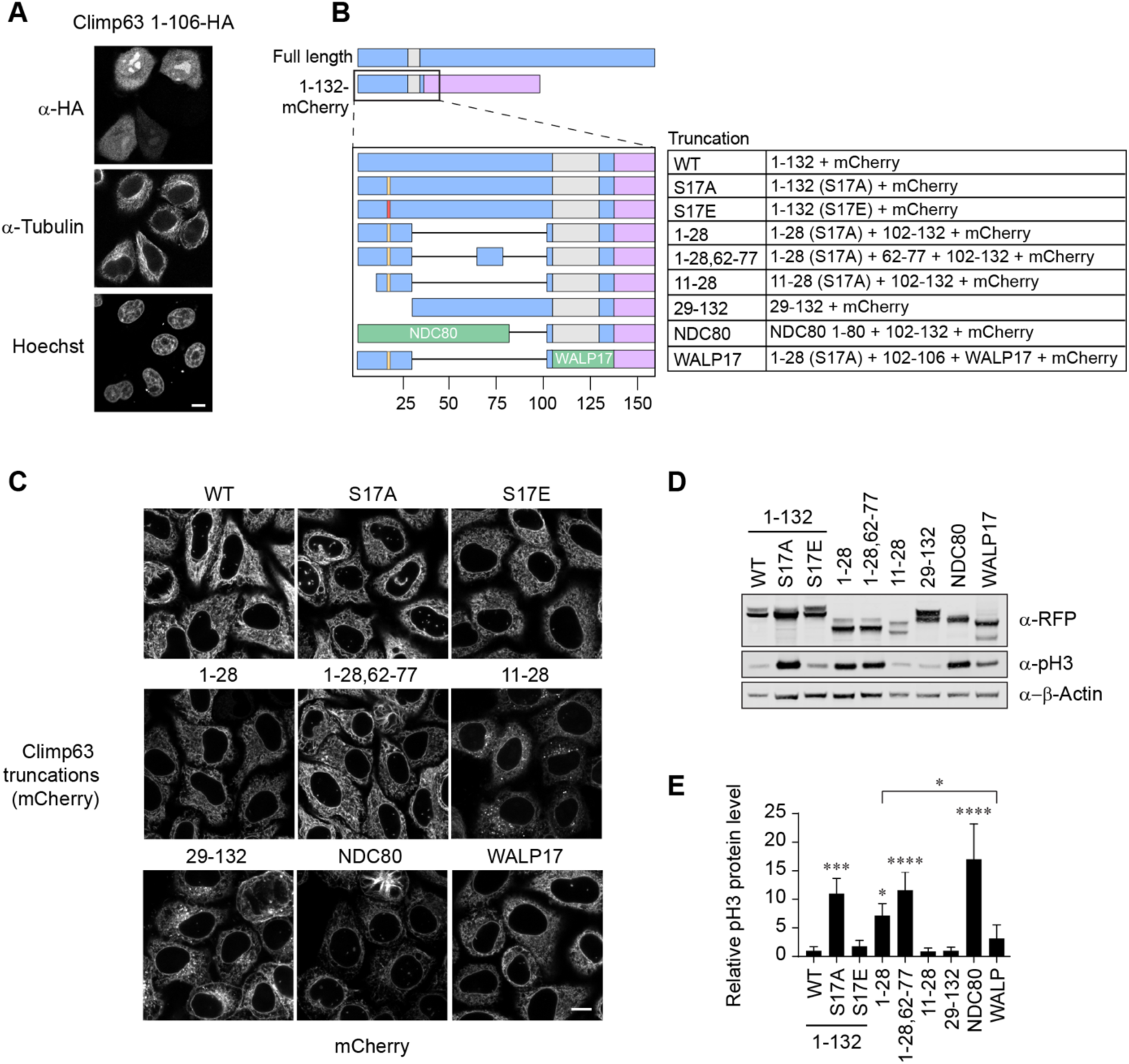
The N-terminal 28 amino acids of the cytoplasmic domain of Climp63 are sufficient for MT-ER interactions. **(A)** Immunofluorescence analysis of the cytoplasmic domain of Climp63 (Climp63 1-106-HA). Scale bar, 10 µm. N = 3. **(B)** Schematic depiction of the generated fragments of the cytoplasmic domain of Climp63. All constructs include a transmembrane domain followed by a C-terminal mCherry. The orange line at position 17 indicates the S17A mutation in these constructs and the red line the S17E mutation.**(C)** Immunofluorescence analysis of tetracycline-inducible HeLa cell lines expressing the depicted Climp63 fragments. Cell lines expressing construct 11-28, 29-132, NDC80 and WALP17 were induced with 1 µg/ml Tetracycline and the other constructs with 20 ng/ml. N = 3. **(D)** Immunoblot analysis of the same samples as in (C) with the indicated antibodies. N = 3. **(E)** Quantification of phospho-H3 (pH3) protein levels in (D) normalized to β-Actin and to the mean within replicates. Asterisks denote significant differences compared to the parental control, unless otherwise indicated. Mean ± SD, *p ≤ 0.05, ***p ≤ 0.001, ****p ≤ 0.0001, N = 4.

Given the lack of binding of the cytoplasmic domain of Climp63 to MTs, we reasoned that the membrane-bound phosphodeficient Climp63 S17A mutant may represent a convenient tool to delineate the regions required for Climp63 MT binding *in vivo* by using the increase in mitotic cells as a readout for Climp63-MT interactions. We therefore generated several truncated fusion constructs, comprising pieces of the cytoplasmic domain of Climp63 containing the S17A mutation, the transmembrane domain and a C-terminal mCherry for visualization (Figure 3B). The first construct contained the N-terminal 28 residues of Climp63 surrounding phosphorylation site S17 (1-28(S17A)). A second construct (1-28,62-77(S17A)) additionally included residues 62-77, as this region has been implicated in MT binding before (Zheng et al., 2022). In addition, we generated a construct lacking the N-terminal 10 residues (11-28(S17A)). In another construct, we exchanged the cytoplasmic domain of Climp63 with the MT binding N-terminal tail of the well-characterized MT binding factor NDC80 (Alushin & Nogales, 2011), which is a kinetochore component. Like Climp63, NDC80 has a basic and unstructured N-terminal tail that binds MTs (Alushin et al., 2012) and we therefore wished to test if the Climp63 and NDC80 unstructured tail regions are interchangeable. Lastly, we exchanged the TM domain of Climp63 for an artificial TM domain (WALP17) to investigate a potential role of its TM domain (de Planque & Killian, 2003), e.g. by facilitating the binding to another membrane-bound component.

Tetracycline-inducible cell lines harboring these constructs were analyzed by confocal microscopy, which showed that all constructs localized to the ER as expected (Figure 3C). To examine MT binding, we harvested cells and analyzed the accumulation of mitotic cells by immunoblotting of phospho-H3 (S10). We observed an increase in mitotic cells for the cell lines expressing Climp63 1-132(S17A), 1-28(S17A), 1-28,62-77(S17A) and the NDC80-Climp63 hybrid (Figure 3C, D), indicating a persistent interaction of these fusion proteins and thereby the ER with MTs in mitosis. The construct harboring the truncated N-terminal region (11-28(S17A)) did not result in the accumulation of mitotic cells as deduced from the unchanged phospho-H3 signal, suggesting that this shortened fragment does not bind to MTs efficiently. We further observed that the phospho-H3 signal caused by the expression of the construct 1-28,62-77(S17A) was higher than that of 1-28(S17A), supporting previous work suggesting an involvement of region 62-77 in MT binding (Zheng et al., 2022). However, although this region might facilitate MT interaction, it seems not sufficient for MT binding, as construct 29-132 does not show an increase in mitotic cells. Collectively, this data shows that the 28 N-terminal amino acids of Climp63 are key for MT binding, with region 62-77 further promoting the association of Climp63 with MTs.

Interestingly, we also observed that exchanging the cytoplasmic domain of Climp63 with residues 1-80 of NDC80 resulted in an increase in mitotic cells similar to Climp63 S17A (Figure 3D, E). This suggests that the NDC80 and Climp63 unstructured tails are functionally interchangeable for MT association in the context of Climp63. In addition, exchanging the TM domain in Climp63 1-28(S17A) to WALP17 reduced MT-ER interactions, indicating that the Climp63 TM domain may promote MT binding. As it is unlikely that this domain would bind to MTs directly, this may suggest that MT binding of Climp63 is facilitated by another protein that interacts with Climp63 via its TM domain. Mechanistically, this could resemble the case of the NDC80 tail, for which binding is supported by its proximal calponin homology (CH) domain (Alushin et al., 2012, Cheeseman et al., 2006, Ciferri et al., 2008, Wei et al., 2007). Altogether, these findings indicate that the N-terminal 28 residues mediate the interaction with MTs, supported by residues 62-77 and the TM domain, and that releasing Climp63-MT interaction is critical for proper mitotic progression and the prevention of mitotic defects.

### Cells expressing phosphodeficient Climp63 arrest in prometaphase due to activation of the spindle assembly checkpoint

To gain a better understanding of how mitosis is affected by the expression of phosphodeficient Climp63, we followed Climp63 WT, 3A and 3E expressing cells through mitosis by time-lapse imaging with the ER marker Sec61β-GFP and the DNA marker SiR-Hoechst. We performed automated image analysis using the CellCognition software to track individual cells through mitosis and annotate their mitotic stage based on chromatin morphology (Held et al., 2010). From this analysis we obtained cell trajectories of mitotic events ranging from 30 min before the prophase to prometaphase transition until 150 min after. Plotting these cell trajectories showed a clear prolongation of prometaphase upon Climp63 3A expression, but not for Climp63 WT and 3E (Figure 4A). Indeed, quantification showed a strong increase of prometaphase duration for Climp63 3A expressing cells (Figure 4B), although the exact mean time in prometaphase could not be determined since the extraordinarily long prometaphase frequently exceeded the total time of the recorded cell trajectories.

**Figure 4.**
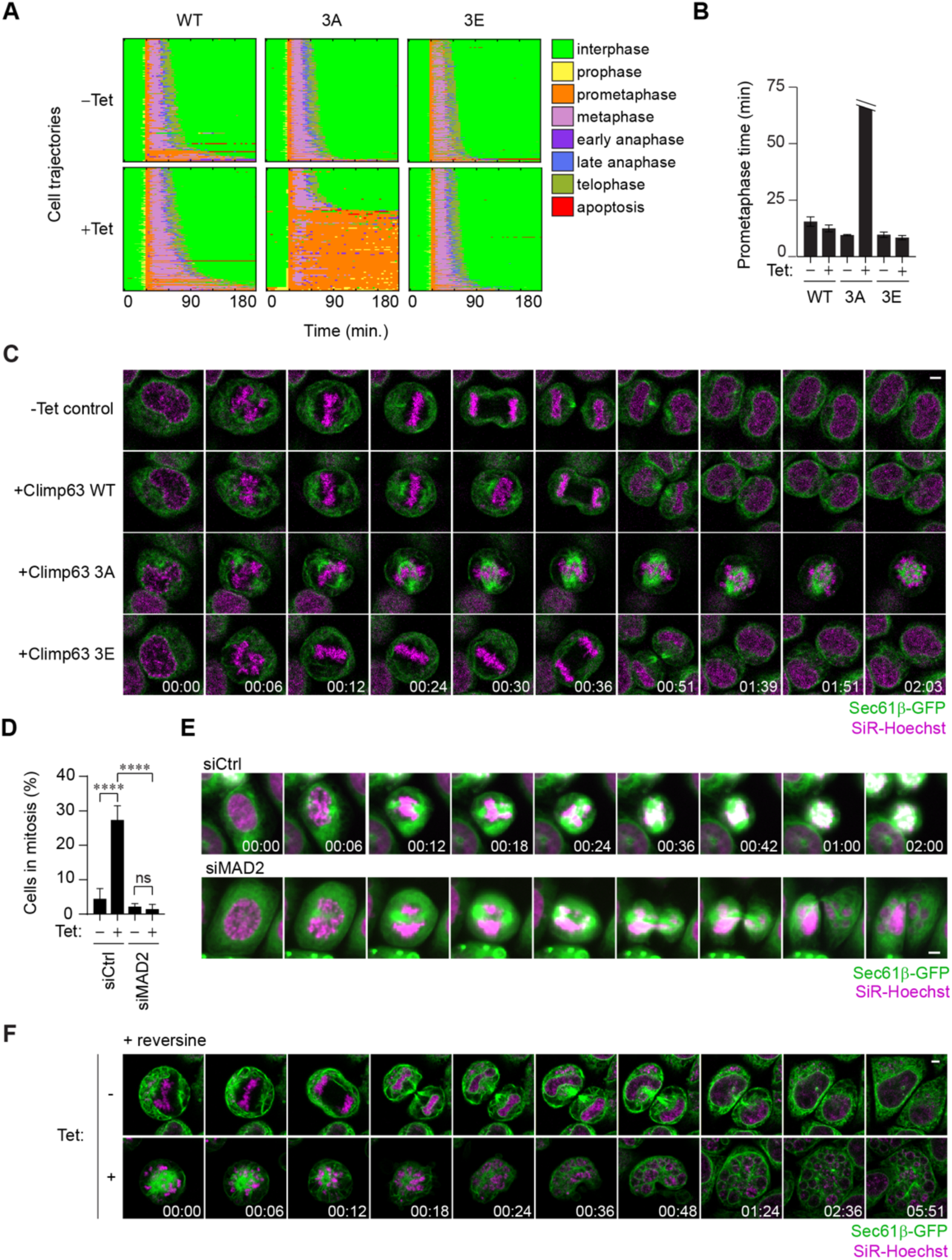
Climp63 3A expressing cells arrest in mitosis due to active SAC signaling. **(A)** Automated image analysis using CellCognition was used to extract mitotic timing from time-lapse imaging of HeLa cell lines expressing Climp63 WT, 3A or 3E for 24 h and imaged every 3 min for a total of 16 h. Cell trajectories (n ≥ 90) are aligned to the prophase to prometaphase transition and color-coded based on the depicted mitotic phases or cell state. **(B)** Quantification of prometaphase length from three replicates as in (A). The maximum length is capped off as trajectories are tracked for 150 min after prophase to prometaphase transition, so the full duration of prometaphase in Climp63 3A-expressing cells is unknown. N = 3, n ≥ 400. **(C)** High resolution time-lapse images of Climp63 WT, 3A or 3E expressing and control HeLa cells progressing through mitosis as in (A). DNA was visualized by SiR-Hoechst and the ER by the stable expression of Sec61β-GFP. Scale bar, 5 µm. **(D)** Quantification of cells in mitosis upon depletion of MAD2 compared to the control. N = 3, n ≥ 800. **(E)** Time-lapse images of a representative cell undergoing mitosis upon 20 nM MAD2 siRNA for 48 h or control treatment. Expression of Climp63 3A was induced for 24 h. Scale bar, 5 µm. **(F)** Representative time-lapse images of mitotic exit in Climp63 3A mutant cells and control upon 320 nM reversine treatment. Reversine was added approximately 10 min before timepoint 0. Scale bar, 5 µm.

Next, higher resolution imaging was used to more closely inspect mitotic progression. Upon mitotic entry, chromosomes congressed to the metaphase plate in all conditions within ∼ 12 min after the initiation of prophase (Figure 4C). At this stage, the ER was observed to progressively infiltrate the spindle area in Climp63 3A expressing cells and eventually completely colocalized with the mitotic spindle (Figure 4–figure supplement 1). Whereas anaphase was initiated for all other conditions within 40 min, Climp63 3A cells failed to enter anaphase. As time progressed, the initially aligned chromosomes started to scatter in Climp63 3A-expressing cells.

The failure to initiate anaphase suggests that the spindle assembly checkpoint (SAC) is active in these cells. Indeed, depletion of the SAC factor MAD2 drastically reduced the number of mitotic cells upon the expression of Climp63 3A (Figure 4D), and cells exited mitosis in a normal time window, albeit accompanied by severe nuclear fragmentation (Figure 4E). Taken together, this data suggests that ER attachment to the mitotic spindle does not impair initial chromosome congression but prevents stable attachment of the spindle to kinetochores, which is required to initiate anaphase.

To visualize how mitotic exit of Climp63 3A expressing cells leads to nuclear fragmentation at high resolution, we induced mitotic exit by overriding the SAC with the Mps1 inhibitor reversine and performed confocal time-lapse microscopy. At the start of imaging, chromosomes in Climp63 3A expressing cells were scattered and the ER was aligned along with what looked like monopolar spindles (Figure 4F). When anaphase was initiated following reversine treatment, chromosomes were not segregated into daughter cells and the cleavage furrow failed to form in most cells, unlike in control cells that have a defined bipolar spindle. Instead, chromosomes remained stationary and spatially separated. Afterwards, chromosomes slowly decondensed and assembled a NE from adjacent ER membranes, resulting in cells filled with many micronuclei. This suggests that the spatial separation of chromosomes by the mislocalized ER is incompatible with chromosome clustering mechanisms, which involve factors such as BAF and Ki67 (Cuylen-Haering et al., 2020, Samwer et al., 2017). Instead, once NE reformation is completed, fragmented nuclei remain physically separated by NE/ER membranes.

From the time-lapse experiments it became evident that ER attachment to the spindle eventually causes it to collapse into a seemingly monopolar configuration. To gain a better understanding of how persistent Climp63-mediated ER-MT interactions affect the spindle, we set out to analyze spindle morphology with and without the expression of Climp63 3A. We chose to investigate cells with aligned metaphase plates for both conditions, with the aim to compare spindles from cells that were in mitosis for a similar time (see Figure 4C). Analysis of z-stacks capturing the entire mitotic spindle showed a relatively normal spindle morphology for Climp63 3A expressing cells at this early time point, although spindles appear slightly more irregular and have a higher eccentricity than control spindles (Figure 5A, B). This suggests that the initial formation of the spindle is not greatly affected by persistent MT-ER contacts.

**Figure 5.**
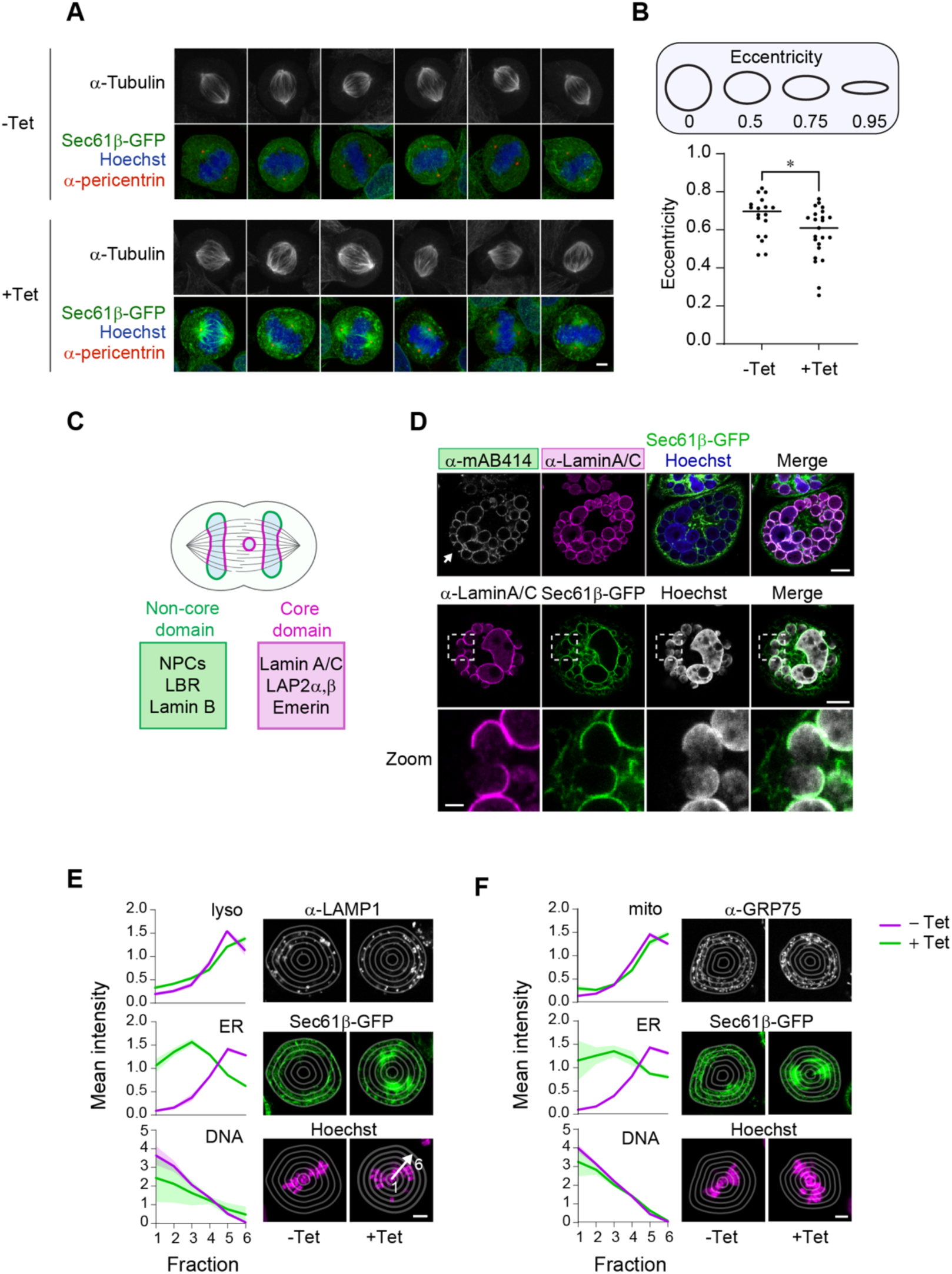
Mitotic consequences of persistent MT-ER interactions induced by the expression of phosphodeficient Climp63. **(A)** Immunofluorescence analysis of spindle morphology of cells with and without tetracycline-induced expression of Climp63 3A. Images are the maximum projections of z-stacks. Scale bar, 5 µm. **(B)** Quantification of tubulin morphology from 3D stacks with and without tetracycline-induced expression of Climp63 3A as in (A). Only cells with mostly aligned chromosomes were quantified, as these have likely not been in mitosis for long yet (see Figure 4C). Horizontal line represents the mean. N = 3, n ≥ 18. *p ≤ 0.05. **(C)** Schematic depiction of NE assembly on a telophase cell. Core components (pink) reassemble in regions with a high MT density, whereas non-core components (green) assemble in regions with low MT density. Lagging chromosomes (represented by the circle) are frequently surrounded by MTs and then assemble only core components, explaining the low NPC density often found on micronuclei formed from lagging chromosomes. **(D)** Immunofluorescence analysis of fragmented nuclei after 72 h of Climp63 3A expression stained for the indicated antibodies that represent core (Lamin A/C; pink box) and non-core (mAB414; green box) proteins. The white arrow points to a part of the nuclear envelope of a micronucleus that is seemingly missing mAB414 and Lamin A/C signal. The bottom image shows this more clearly with a zoom-in. Upper two scale bars, 10 µm, scale bar of zoom-in, 2 µm, N = 3. **(E, F)** Representative immunofluorescence images and quantification of the radial distribution of lysosomes (E, α-LAMP1) and mitochondria (F, α-GRP75), which were analyzed in 6 bins using CellProfiler, with and without expression of Climp63 3A. The lines represent the mean intensity per bin, shaded areas ± SD. N = 3, n > 18. Scale bar, 5 µm.

### The NE assembles properly on fragmented nuclei

Next, we examined the consequences of nuclear fragmentation on NE reformation by investigating the composition of the NE surrounding the formed micronuclei. The composition of the NE surrounding micronuclei is critical, because a failure to incorporate a subset of NE proteins makes them prone to rupture. NE rupture in turn can result in chromothripsis (Leibowitz et al., 2015), a process in which a genomic region is shattered and stitched together randomly, accompanied by a high mutation rate that is disastrous for genomic integrity and has been observed in up to 50% of adult human cancers (Cortes-Ciriano et al., 2020, Simovic-Lorenz & Ernst, 2025, Voronina et al., 2020). Fragile micronuclei particularly lack the so-called non-core NE proteins (Liu et al., 2018). Non-core NE proteins are characterized by their assembly on the chromatin surface away from the central spindle, whereas core proteins enrich adjacent to the spindle (Figure 5C) (Haraguchi et al., 2008). Lagging chromosomes are frequently surrounded by MTs and are therefore thought to lack non-core components (Liu et al., 2018). Nuclear pore complexes (NPCs) belong to the non-core group and are therefore frequently absent from micronuclei formed around lagging chromosomes. A lack of NPCs renders these micronuclei incompetent to import the factors required for the maintenance of genome integrity (Liu et al., 2018).

To gain insights into the composition of the NE surrounding the micronuclei formed upon expression of phosphodeficient Climp63 3A, we performed immunofluorescence analysis of FG nucleoporins (mAB414), a readout for non-core proteins, and lamin A/C, a core protein (Figure 5D). The fragmented nuclei contained both FG nucleoporins and lamin A/C with a relatively equal distribution among all micronuclei, showing that both groups of NE proteins were recruited. This indicates that the mode of NE assembly on these fragmented nuclei is different from that usually observed on lagging chromosomes (Liu et al., 2018, Zhao et al., 2023) and suggests that they might be more stable than micronuclei that lack a subset of NE proteins. Consistently, we did not observe any features indicative of micronuclear rupture such as compacted chromatin, accumulation of Sec61β-GFP and the absence of NPCs (Ferrandiz et al., 2022, Hatch et al., 2013), although we cannot exclude that these micronuclei may rupture transiently. In addition, time-lapse imaging revealed that cells survive for at least 6 h with a fragmented nucleus (see Figure 4F). Interestingly, around individual micronuclei, the distribution of NE components is not uniform, as some regions seem devoid of Sec61β-GFP or NE/ER membrane components. While we cannot rule out imaging artefacts, we have consistently observed this throughout z-stacks and for all studied marker proteins. Still, altogether our data shows that Climp63-mediated nuclear fragmentation does not significantly impair NE reformation.

### Repositioning of the ER does not affect mitochondrial or lysosomal positioning in mitosis

In interphase, the ER affects the positioning and division of various organelles (Prinz et al., 2020). Although not much is known about remodelling of ER-organellar contact sites in mitosis (Kors & Schlaitz, 2024), it is established that mitochondria are closely associated with the ER during interphase and these contact sites are thought to expand during mitosis (Moore et al., 2021, Yu et al., 2024). The repositioning of the ER to the mitotic spindle upon Climp63 3A expression provides a simple way to investigate how the ER affects the positioning of other organelles in mitosis.

To investigate whether repositioning of the ER upon Climp63 3A expression affects other organelles, we performed immunofluorescence staining to visualize mitochondria (GRP75), lysosomes (LAMP1), ER (Sec61β), and DNA and analyzed how their radial distribution was affected by the expression of Climp63 3A. For the quantifications, we measured the distribution of the chosen markers in a series of six radial bins. As a proof of principle, the quantification pipeline showed that the mean intensity of DNA was the highest in the most central radial bin and decreased in intensity peripherally (Figure 5E, F). In absence of Climp63 3A expression, the ER follows the opposite distribution, with a higher intensity in the cell periphery. Expression of Climp63 3A repositions the ER to the spindle area, resulting in a higher mean intensity in the central bins. Interestingly, the expression of Climp63 3A did not affect the radial distribution of mitochondria or lysosomes (Figure 5E, F), suggesting that these organelles do not follow the ER in their mitotic positioning. This indicates that the positioning of the organelles is determined by other mechanisms, independently of the ER.

## Discussion

Upon mitotic entry, most connections between the ER and the MT network are broken to allow for the separation of membranes from the mitotic spindle and chromatin (Champion et al., 2019, Kors & Schlaitz, 2024). Our work demonstrates the importance of breaking the connections between ER sheets and MTs for the example of the MT binding protein Climp63, which is highly abundant in ER sheets. Expression of a Climp63 phosphorylation-deficient mutant (S3A, S17A, S19A) hinders mitotic progression and leads to severe nuclear fragmentation upon mitotic exit, which can result in the formation of up to 30 or more micronuclei in one cell. To our knowledge, such striking nuclear fragmentation as a consequence of a mitotic error has not been reported before. One comparable case of nuclear fragmentation occurs when mitotic exit is induced in taxol-treated cells (Samwer et al., 2017). This fragments the nucleus, likely because the taxol-mediated MT stabilization results in the forced separation of chromosomes as the NE reforms, similar to what we observe upon expression of phosphorylation-deficient Climp63.

### MT binding of Climp63

The fact that phosphorylation-deficient Climp63 causes a stark increase in mitotic cells because of persistent MT-ER interactions allowed us to study the MT binding ability of Climp63, using the number of mitotic cells as a readout. Previous studies have investigated the binding of Climp63 to MTs by co-sedimentation assays with varying results (Klopfenstein et al., 1998, Vedrenne et al., 2005, Zheng et al., 2022). Since co-sedimentation assays can be subject to non-specific binding, especially in cell lysates, our system provides a useful *in vivo* approach. We identify a key role of the N-terminal region (residues 1-28) of Climp63 in MT binding, with an additional contribution of residues 62-77, in agreement with a previous study (Zheng et al., 2022).

The minimal region we found to bind MTs (aa 1-28) is predicted to be unstructured and basic (pI = 10.3). Although many MT binding proteins possess structured domains to bind to MTs, such as a CH domain, CAP-Gly domains, and TOG domains (Ayaz et al., 2012, Slep & Vale, 2007, Wang et al., 2014), other MT binding proteins exploit unstructured, basic domains instead, including the inner kinetochore protein CENP-Q, the Alzheimer’s disease-related Tau and the kinetochore protein NDC80 (El Mammeri et al., 2022, Pesenti et al., 2018, Weingarten et al., 1975). In our experiments, we demonstrated that the N-terminal tail of NDC80 can functionally replace the Climp63 cytoplasmic domain, confirming their functional and structural similarity.

Furthermore, we explored the individual contribution of the three phosphorylation sites that were initially suggested to regulate MT binding of Climp63 (Vedrenne et al., 2005). We identified S17 as the key residue to impair MT binding upon its phosphorylation and we show that this site is phosphorylated by the master regulator of mitosis, CDK1. As the other two sites do not seem to affect MT binding, these sites might instead regulate Climp63 function and activity in other settings. One hypothesis for their function comes from a study that implicates both PKC and Climp63 in controlling the rate-limiting step of cargo transport to the plasma membrane in neuronal dendrites (Wang et al., 2012). Climp63 was suggested to increase dendritic branching by PKC-mediated phosphorylation of S3, S17 or S19 or a combination thereof, and thereby confine cargo in the neuronal dendrites (Wang et al., 2012). Given that site S3 and S19 might be PKC targets (Vedrenne et al., 2005), this data suggests that these sites could affect ER architecture and dendritic branching, while our data shows that S17 regulates the affinity for MTs and is a target of CDK1 instead.

An important remaining question about the binding of Climp63 to MTs is whether it is direct or indirect. We observed that the cytoplasmic domain of Climp63 (1-106) does not colocalize with MTs, in contrast to a previous report (Klopfenstein et al., 1998). Our data instead indicate that membrane association of Climp63 may provide the avidity needed to bind MTs or that the interaction is indirect and facilitated by another (membrane-bound) component. In support of the latter, we find that the TM domain of Climp63 promotes MT binding, which might indicate the involvement of a binding partner. Furthermore, the existing data for direct binding lack appropriate negative controls, making it hard to exclude unspecific binding in the *in vitro* assay (Klopfenstein et al., 1998). Overall, future experiments will need to further dissect how the N-terminal 28 amino acids together with aa 62-77 of Climp63 bind to MTs, what binding partners might facilitate its MT binding, and how the Climp63 TM domain may contribute.

### How does ER attachment to the mitotic spindle affect mitosis?

In our experiments, we observed a drastic delay in mitotic progression upon the expression of phosphodeficient Climp63. Affected cells were arrested in prometaphase, the stage in which chromosomes congress to the metaphase plate. Interestingly, high resolution time-lapse imaging of mitotic onset showed that chromosomes initially line up at the metaphase plate, but scatter over time when anaphase is not initiated. So how does ER attachment to the mitotic spindle affect mitotic processes? The fact that chromosomes line up initially suggests that motor-based movements and MT (de)polymerization, both required to direct chromosomes to the metaphase plate, are not impaired. However, cells arrested in prometaphase can be experimentally induced to progress through mitosis by inhibition of the SAC machinery, indicating that the SAC remains active upon Climp63-mediated ER tethering to MTs. There are several ways in which persistent binding of Climp63 to MTs could interfere with silencing of the SAC. First, ER membranes may physically block MTs, preventing them from forming stable attachments at kinetochores. Second, the mitotic checkpoint complex (MCC) is usually actively removed from kinetochores by dynein/dynactin to silence the SAC (Griffis et al., 2007, Howell et al., 2001, McAinsh & Kops, 2023, Wojcik et al., 2001). Possibly, ER attachment to the mitotic spindle might impair this dynein-dependent stripping of the MCC from kinetochores and thereby halt anaphase progression by continued inhibition of the anaphase-promoting complex (APC/C). Finally, satisfaction of the SAC relies on a conformational change in NDC80 upon binding to MTs, which releases the Mps1 kinase from kinetochores to facilitate anaphase initiation (Hiruma et al., 2015, Ji et al., 2015). Given the similarities between the unstructured N-terminal tails of Climp63 and NDC80, Climp63 could occupy the same binding sites on MTs as NDC80 and compete with its binding, thereby preventing the stable attachment of NDC80 with MTs, if these binding sites were at all limiting. These models might explain how anaphase progression is impaired by Climp63-mediated persistent ER-MT interactions.

We further demonstrated that persistent mitotic ER-MT interactions resulted in the formation of many micronuclei during mitotic exit. Micronuclei are often fragile and prone to chromothripsis (Hatch et al., 2013, Liu et al., 2018, Maciejowski & Hatch, 2020), largely due to compromised NE integrity caused by the absence of NPCs and non-core proteins on micronuclei. A number of studies set out to investigate why NPCs are frequently excluded from micronuclei, resulting in several hypotheses: proximal MTs might block NPC assembly, membranes assembling on lagging chromosomes might lack the topology required to template NPCs, and delayed membrane recruitment on lagging chromosomes might hinder their NPC assembly (Liu et al., 2018, Zhao et al., 2023). Interestingly, we found that all micronuclei induced by the persistent interaction of Climp63 with MTs contain NPCs. This suggests that MT proximity may not affect NE assembly *per se*, and that lagging chromosomes might lack NPCs and non-core proteins for a different reason.

Overall, our findings highlight the importance of regulating MT-ER interactions in mitosis to allow for the formation of a single nucleus and provides a new perspective on the growing understanding of the role of the ER in mitosis.

## Materials and Methods

### Antibodies

Commercial antibodies used were anti-Climp63 (16686-1-AP, rabbit, Proteintech), anti-phospho histone H3 (S10) (9701S, rabbit, Cell Signaling Technology), anti-β-actin (sc-47778, mouse, Santa Cruz), anti-tubulin (T5168, mouse, Sigma Aldrich), anti-mAB414 (ab24609, mouse, Abcam), anti-lamin A/C (10298-1-AP, rabbit, Proteintech), anti-LAMP1 (sc-20011, mouse, Santa Cruz), anti-GRP75 (ab2799, mouse, Abcam), anti-pericentrin (ab4448, rabbit, Abcam).

### Molecular cloning

A plasmid containing the corresponding cDNA fragment for Climp63 was a gift from Peter Steyger (Addgene plasmid # 80977) and was used to clone Climp63 into the pcDNA5 FRT/TO vector (CMV promotor, Invitrogen) for the generation of stable tetracycline-inducible cell lines. The NDC80 coding region was amplified from HeLa cDNA and cloned into pcDNA5 FRT/TO. Climp63 served as a PCR template to generate Climp63 1-106 and 1-132, which were cloned into pcDNA5-HA FRT/TO and pcDNA5-mCherry FRT/TO, respectively. Truncations of Climp63 1-132 and HisSUMO-PreScission-Climp63-1-32-GFP-HA for E. coli expression were obtained from Twist Bioscience and were codon-optimized and cloned into pcDNA5-mCherry FRT/TO and the pQE30 expression vector, respectively. To generate single and triple mutants (S3, S17, S19 to A or E), full-length Climp63, Climp63 1-132, and HisSUMO-PreScission-Climp63-1-31-GFP-HA were used as templates for mutagenesis with the QuikChange mutagenesis kit (Agilent Technologies). The Climp63-WALP17 hybrid was cloned using the WALP17-mCherry template from (Ungricht et al., 2015). In addition, the ER targeting of the WALP17 construct and Climp63 11-28 was optimized by including an RRSR targeting motif right after amino acid 28, followed by a GSSGGGSG linker before the TM domains.

### Cell culture and cell lines

HeLa cells were grown in Dulbecco’s modified Eagle medium (DMEM) supplemented with 10% fetal calf serum and 100 μg/ml penicillin/streptomycin at 37°C with 5% CO_2_. Stable tetracycline-inducible cell lines were generated by integration into FRT sites and were selected with 0.4 mg/ml hygromycin B. Expression of Climp63 was induced with 50 ng/ml tetracycline (Sigma-Aldrich) for all experiments unless indicated otherwise. The HeLa cell line stably expressing Sec61β-GFP has been described previously (Pawar et al., 2017). Transfections of 20 nM Allstars control (Qiagen) and MAD2 (ATGGATATTTGTACTGTTTAA; Qiagen) siRNA oligonucleotides were performed with the Lipofectamine RNAiMax (Invitrogen) transfection reagent. siRNAs were incubated for 48 h before imaging.

### Immunofluorescence analysis and confocal microscopy

Cells were grown on glass coverslips and fixed in 4% paraformaldehyde in PBS for 10 min. Cells were permeabilized with 0.2% Triton X-100 in PBS for 5 min and blocked with 2% BSA/PBS for 30 min. For lysosomal stainings, cells were fixed in 3.7% formaldehyde and permeabilized and washed with 10% BSA, 15 mM glycine, 0.05% saponin and 10 mM HEPES. Primary antibodies, diluted in 2% BSA/PBS, were incubated for 1 hr. After three washes with 2% BSA/PBS, secondary antibodies and Hoechst in 2% BSA/PBS were incubated for 30 min After three washes, coverslips were mounted onto microscopic slides with VectaShield (VectorLabs). All steps were performed at RT. Imaging was performed with a confocal microscope (Zeiss LSM780, 63X 1.4 NA oil DIC Plan-Apochromat objective).

### Live cell imaging

Cells were seeded on 8-well LabTek chambers the day before imaging. Two hours before live imaging, culture medium was changed to FluoroBrite DMEM supplemented with 10% fetal calf serum and 100 μg/ml penicillin/streptomycin and either 0.1 mM SiR-Hoechst (Spirochrome), 1:1000 SPY650-DNA (Spirochrome), and 1:1000 SPY555-tubulin (Spirochrome) as indicated for each experiment. All live cell imaging experiments were imaged on a Zeiss Celldiscoverer 7 in a 5% CO_2_, 37°C chamber.

For the time-lapse microscopy of mitotic exit upon reversine treatment a 2D Airyscan super-resolution/SR mode was used. Images were taken every 3 min for 6 h with 12 z-stacks using a 20x 0.95 NA Plan-Apochromat objective with a 2x tubelens (40x total). Images were analyzed with the Airyscan Processing function in Zen Black software with the default settings.

Imaging of mitotic entry was performed using a 20x 0.95 NA Plan-Apochromat objective with a 2x tubelens (40x total). Images were taken every 3 min for 6 h with 10 z-stacks. For the mitotic timing analyses, images were taken every 3 min for 10 h with a Plan-Apochromat 5x/0.35 objective and 2x tubelens (10x total). For the MAD2 RNAi time-lapse imaging, cells were imaged every 3 min for 5 h with 3 z-stacks with a Plan-Apochromat 20x/0.7 objective.

### Immunoblot analysis

Cells were resuspended in SDS sample buffer (75 mM Tris pH 6.8, 20% (v/v) glycerol, 4% (w/v) SDS, 50 mM DTT, 0.01% (w/v) bromophenol blue), sonicated and boiled at 95°C for 5 min. Samples were separated on SDS-PAGE gels and transferred onto a nitrocellulose membrane using semi-dry blotting. Membranes were blocked in 5% milk (w/v) in PBST (0.1% Tween-20 in PBS) and incubated with primary antibodies in 4% milk/PBST overnight. After three washes for 5 min with PBST, secondary antibodies (goat anti-mouse Alexa Fluor Plus 680 (Thermo Fisher Scientific, A32729) or goat anti-rabbit Alexa Fluor Plus 800 (Thermo Fisher Scientific, A32735) in 4% milk/PBST were incubated for at least 30 min at RT. After three final washes with PBST, blots were imaged and quantified using an Odyssey (LI-COR) imaging system.

### Image analysis

Mitotic timing was analyzed with the CellCognition software v1.6.1 (Held et al., 2010). Chromatin morphology visualized by SiR-Hoechst was used to classify mitotic stages as described in (Held et al., 2010). Segmentation was achieved with a local adaptive threshold and cells were tracked using a constrained nearest-neighbor approach with trajectory splitting and merging. Cell trajectories were selected based on the detection of a prophase to prometaphase transition and included 30 min before and 150 min after this transition.

For organelle positioning experiments, the mid-plane of a mitotic cell with mostly aligned chromosomes was imaged and quantified with CellProfiler. Cell boundaries were segmented based on the Sec61β-GFP signal. Cells were radially divided into six bins and the mean intensity of each bin was calculated with the MeasureObjectIntensityDistribution module. The mean for each bin and replicate was plotted in GraphPad Prism.

For imaging spindle morphology, z-stacks were taken that capture the entire cell. Maximum z-projections were analyzed in CellProfiler, where cell boundaries were segmented based on the Sec61β-GFP signal. Using the MeasureObjectSizeShape, the eccentricity of the spindle was calculated. The mean eccentricity of each replicate was calculated and plotted with GraphPad Prism.

### Protein purification

6xHis-SUMO-PreScission-Climp63-1-31-GFP-HA WT and S17E were expressed in *E. coli* BLR(pREP4) cells with 0.5 mM IPTG for 5 h at 25°C. Cells were pelleted and lysed in 50 mM Tris, 700 mM NaCl, 1 mM MgCl_2_, 50 mM imidazole, 5 µg/ml DNase and 1 mM β-mercaptoethanol using an EmulsiFlex-C5 high pressure homogenizer (Avestin). The lysate was cleared by centrifugation at 45000 rpm in a Ti70 rotor. The cleared lysate was incubated on a 5 ml HisTrap FF Crude histidine-tagged protein purification column (Cytiva) and washed with lysis buffer. Protein was eluted by increasing concentrations of elution buffer consisting of 50 mM Tris, 700 mM NaCl, 1 mM MgCl_2_, and 400 mM imidazole using an ÄKTA prime system. N-terminal His-SUMO was cleaved off with 3C protease while dialyzing at 4°C overnight in 50 mM HEPES-KOH, 150 mM NaCl, 2 mM MgCl_2_, 1 mM DTT and 5% glycerol. Cleavage products were cleaned up by incubating on a HisTrap FF column and collecting the flow-through.

### Kinase assay

Protein phosphorylation assays were performed with 10 µg of Climp63 1-31-GFP-HA, 1 µg of Histone H1 (Sigma-Aldrich, H1917) and 350 ng cyclinB1-CDK1 (Thermo Fisher Scientific, PV3292) in kinase buffer containing 100 µM cold ATP and 2.5 μCi ^32^P-γ-ATP, 20 mM HEPES-KOH, 110 mM KOAc, 5 mM MgCl_2_ and 1 mM DTT. Reactions were incubated for 15 min at 30°C and inactivated by the addition of SDS sample buffer. Samples were analyzed by SDS-PAGE and autoradiography.

## Acknowledgements

We thank H. Baird, V. Sinigiani and I. Zemp for critical reading of the manuscript and other members of the Kutay lab for helpful discussions. Microscopy was performed on instruments of the ETHZ Microscopy Center (ScopeM). We thank F. Marxer, M. Wieczorek, R. Klemm, B. Rowland, and J. Corn for their input. This work was supported by the Swiss National Science Foundation (310030_184801 to U.K.)

---

**Figure 4–figure supplement 1.**
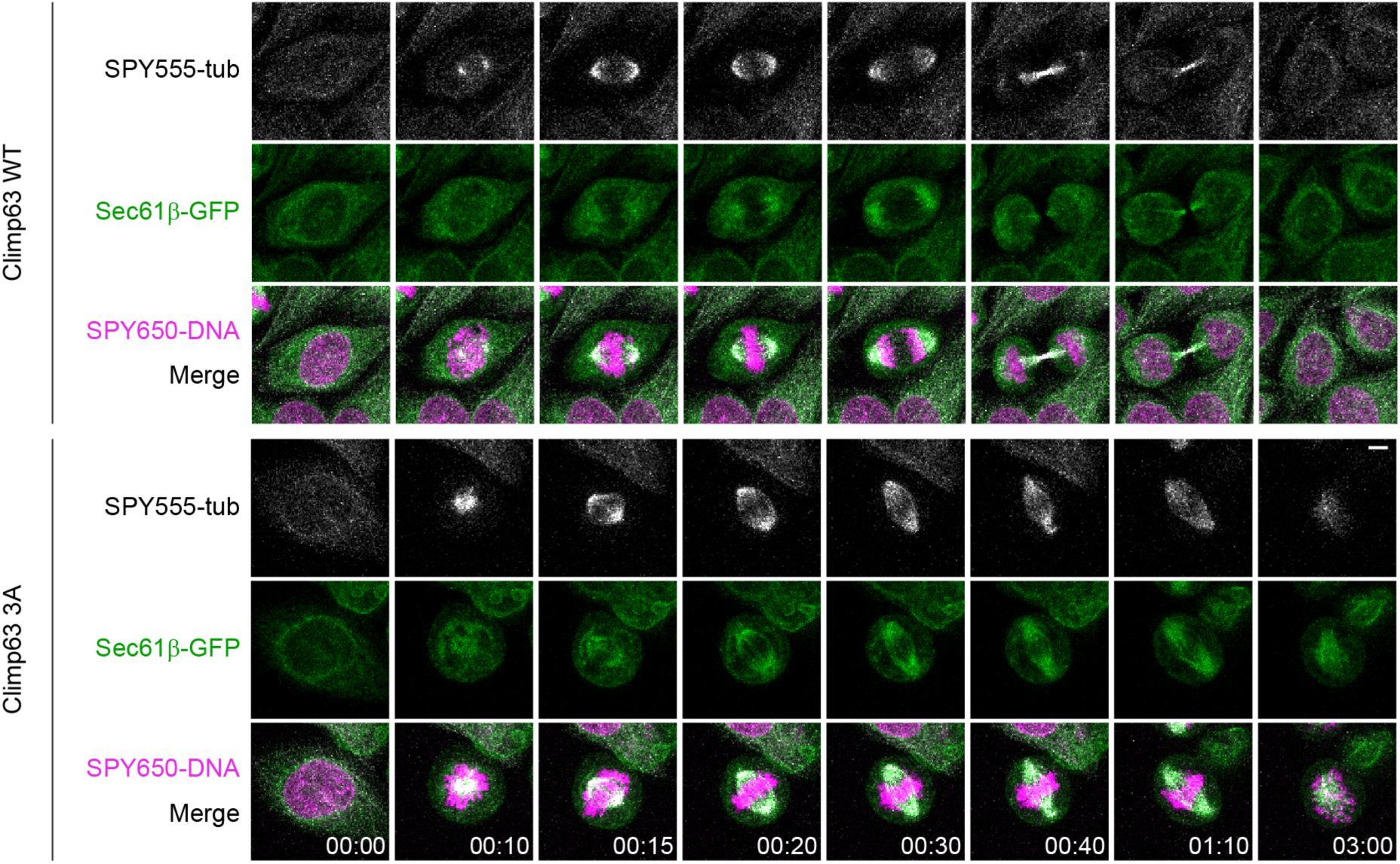
Combined time-lapse imaging of tubulin and ER in Climp63 WT and 3A expressing cells. Representative time-lapse images of Climp63 WT and 3A-expressing HeLa cells progressing through mitosis. DNA was visualized by SPY650-DNA and tubulin by SPY555-tubulin. Scale bar, 5 µm.

## References

Alushin G, Nogales E (2011) Visualizing kinetochore architecture. Curr Opin Struct Biol 21: 661–9

Alushin GM, Musinipally V, Matson D, Tooley J, Stukenberg PT, Nogales E (2012) Multimodal microtubule binding by the Ndc80 kinetochore complex. Nat Struct Mol Biol 19: 1161–7

Ayaz P, Ye X, Huddleston P, Brautigam CA, Rice LM (2012) A TOG:alphabeta-tubulin complex structure reveals conformation-based mechanisms for a microtubule polymerase. Science 337: 857–60

Beaudouin J, Gerlich D, Daigle N, Eils R, Ellenberg J (2002) Nuclear envelope breakdown proceeds by microtubule-induced tearing of the lamina. Cell 108: 83–96

Champion L, Pawar S, Luithle N, Ungricht R, Kutay U (2019) Dissociation of membrane-chromatin contacts is required for proper chromosome segregation in mitosis. Mol Biol Cell 30: 427–440

Cheeseman IM, Chappie JS, Wilson-Kubalek EM, Desai A (2006) The conserved KMN network constitutes the core microtubule-binding site of the kinetochore. Cell 127: 983–97

Chung JY, Steen JA, Schwarz TL (2016) Phosphorylation-Induced Motor Shedding Is Required at Mitosis for Proper Distribution and Passive Inheritance of Mitochondria. Cell Rep 16: 2142–2155

Ciferri C, Pasqualato S, Screpanti E, Varetti G, Santaguida S, Dos Reis G, Maiolica A, Polka J, De Luca JG, De Wulf P et al. (2008) Implications for kinetochore-microtubule attachment from the structure of an engineered Ndc80 complex. Cell 133: 427–39

Cortes-Ciriano I, Lee JJ, Xi R, Jain D, Jung YL, Yang L, Gordenin D, Klimczak LJ, Zhang CZ, Pellman DS et al. (2020) Comprehensive analysis of chromothripsis in 2,658 human cancers using whole-genome sequencing. Nat Genet 52: 331–341

Cuylen-Haering S, Petrovic M, Hernandez-Armendariz A, Schneider MWG, Samwer M, Blaukopf C, Holt LJ, Gerlich DW (2020) Chromosome clustering by Ki-67 excludes cytoplasm during nuclear assembly. Nature 587: 285–290

de Planque MR, Killian JA (2003) Protein-lipid interactions studied with designed transmembrane peptides: role of hydrophobic matching and interfacial anchoring. Mol Membr Biol 20: 271–84

Deolal P, Scholz J, Ren K, Bragulat-Teixidor H, Otsuka S (2024) Sculpting nuclear envelope identity from the endoplasmic reticulum during the cell cycle. Nucleus 15: 2299632

El Mammeri N, Dregni AJ, Duan P, Wang HK, Hong M (2022) Microtubule-binding core of the tau protein. Sci Adv 8: eabo4459

Ellenberg J, Siggia ED, Moreira JE, Smith CL, Presley JF, Worman HJ, Lippincott-Schwartz J (1997) Nuclear membrane dynamics and reassembly in living cells: targeting of an inner nuclear membrane protein in interphase and mitosis. J Cell Biol 138: 1193–206

Ferrandiz N, Downie L, Starling GP, Royle SJ (2022) Endomembranes promote chromosome missegregation by ensheathing misaligned chromosomes. J Cell Biol 221: e202203021

Fischle W, Tseng BS, Dormann HL, Ueberheide BM, Garcia BA, Shabanowitz J, Hunt DF, Funabiki H, Allis CD (2005) Regulation of HP1-chromatin binding by histone H3 methylation and phosphorylation. Nature 438: 1116–22

Goyal U, Blackstone C (2013) Untangling the web: mechanisms underlying ER network formation. Biochim Biophys Acta 1833: 2492–8

Griffis ER, Stuurman N, Vale RD (2007) Spindly, a novel protein essential for silencing the spindle assembly checkpoint, recruits dynein to the kinetochore. J Cell Biol 177: 1005–15

Grigoriev I, Gouveia SM, van der Vaart B, Demmers J, Smyth JT, Honnappa S, Splinter D, Steinmetz MO, Putney JW, Jr., Hoogenraad CC et al. (2008) STIM1 is a MT-plus-end-tracking protein involved in remodeling of the ER. Curr Biol 18: 177–82

Guo Y, Li D, Zhang S, Yang Y, Liu JJ, Wang X, Liu C, Milkie DE, Moore RP, Tulu US et al. (2018) Visualizing Intracellular Organelle and Cytoskeletal Interactions at Nanoscale Resolution on Millisecond Timescales. Cell 175: 1430–1442 e17

Haraguchi T, Kojidani T, Koujin T, Shimi T, Osakada H, Mori C, Yamamoto A, Hiraoka Y (2008) Live cell imaging and electron microscopy reveal dynamic processes of BAF-directed nuclear envelope assembly. J Cell Sci 121: 2540–54

Hatch EM, Fischer AH, Deerinck TJ, Hetzer MW (2013) Catastrophic nuclear envelope collapse in cancer cell micronuclei. Cell 154: 47–60

Held M, Schmitz MH, Fischer B, Walter T, Neumann B, Olma MH, Peter M, Ellenberg J, Gerlich DW (2010) CellCognition: time-resolved phenotype annotation in high-throughput live cell imaging. Nat Methods 7: 747–54

Hiruma Y, Sacristan C, Pachis ST, Adamopoulos A, Kuijt T, Ubbink M, von Castelmur E, Perrakis A, Kops GJ (2015) CELL DIVISION CYCLE. Competition between MPS1 and microtubules at kinetochores regulates spindle checkpoint signaling. Science 348: 1264–7

Honnappa S, Gouveia SM, Weisbrich A, Damberger FF, Bhavesh NS, Jawhari H, Grigoriev I, van Rijssel FJ, Buey RM, Lawera A et al. (2009) An EB1-binding motif acts as a microtubule tip localization signal. Cell 138: 366–76

Howell BJ, McEwen BF, Canman JC, Hoffman DB, Farrar EM, Rieder CL, Salmon ED (2001) Cytoplasmic dynein/dynactin drives kinetochore protein transport to the spindle poles and has a role in mitotic spindle checkpoint inactivation. J Cell Biol 155: 1159–72

Itzhak DN, Tyanova S, Cox J, Borner GH (2016) Global, quantitative and dynamic mapping of protein subcellular localization. Elife 5: e16950

Ji Z, Gao H, Yu H (2015) CELL DIVISION CYCLE. Kinetochore attachment sensed by competitive Mps1 and microtubule binding to Ndc80C. Science 348: 1260–4

Klopfenstein DR, Kappeler F, Hauri HP (1998) A novel direct interaction of endoplasmic reticulum with microtubules. EMBO J 17: 6168–77

Kors S, Schlaitz AL (2024) Dynamic remodelling of the endoplasmic reticulum for mitosis. J Cell Sci 137: jcs261444

Leibowitz ML, Zhang CZ, Pellman D (2015) Chromothripsis: A New Mechanism for Rapid Karyotype Evolution. Annu Rev Genet 49: 183–211

Liu S, Kwon M, Mannino M, Yang N, Renda F, Khodjakov A, Pellman D (2018) Nuclear envelope assembly defects link mitotic errors to chromothripsis. Nature 561: 551–555

Maciejowski J, Hatch EM (2020) Nuclear Membrane Rupture and Its Consequences. Annu Rev Cell Dev Biol 36: 85–114

McAinsh AD, Kops G (2023) Principles and dynamics of spindle assembly checkpoint signalling. Nat Rev Mol Cell Biol 24: 543–559

Moore AS, Coscia SM, Simpson CL, Ortega FE, Wait EC, Heddleston JM, Nirschl JJ, Obara CJ, Guedes-Dias P, Boecker CA et al. (2021) Actin cables and comet tails organize mitochondrial networks in mitosis. Nature 591: 659–664

Nikonov AV, Hauri HP, Lauring B, Kreibich G (2007) Climp-63-mediated binding of microtubules to the ER affects the lateral mobility of translocon complexes. J Cell Sci 120: 2248–58

Nourbakhsh K, Ferreccio AA, Bernard MJ, Yadav S (2021) TAOK2 is an ER-localized kinase that catalyzes the dynamic tethering of ER to microtubules. Dev Cell 56: 3321–3333 e5

Obara CJ, Moore AS, Lippincott-Schwartz J (2023) Structural Diversity within the Endoplasmic Reticulum-From the Microscale to the Nanoscale. Cold Spring Harb Perspect Biol 15: a041259

Parlakgul G, Arruda AP, Pang S, Cagampan E, Min N, Guney E, Lee GY, Inouye K, Hess HF, Xu CS et al. (2022) Regulation of liver subcellular architecture controls metabolic homeostasis. Nature 603: 736–742

Pawar S, Ungricht R, Tiefenboeck P, Leroux JC, Kutay U (2017) Efficient protein targeting to the inner nuclear membrane requires Atlastin-dependent maintenance of ER topology. Elife 6: e28202

Pesenti ME, Prumbaum D, Auckland P, Smith CM, Faesen AC, Petrovic A, Erent M, Maffini S, Pentakota S, Weir JR et al. (2018) Reconstitution of a 26-Subunit Human Kinetochore Reveals Cooperative Microtubule Binding by CENP-OPQUR and NDC80. Mol Cell 71: 923–939 e10

Prinz WA, Toulmay A, Balla T (2020) The functional universe of membrane contact sites. Nat Rev Mol Cell Biol 21: 7–24

Salina D, Bodoor K, Eckley DM, Schroer TA, Rattner JB, Burke B (2002) Cytoplasmic dynein as a facilitator of nuclear envelope breakdown. Cell 108: 97–107

Samwer M, Schneider MWG, Hoefler R, Schmalhorst PS, Jude JG, Zuber J, Gerlich DW (2017) DNA Cross-Bridging Shapes a Single Nucleus from a Set of Mitotic Chromosomes. Cell 170: 956–972 e23

Sandoz PA, Denhardt-Eriksson RA, Abrami L, Abriata LA, Spreemann G, Maclachlan C, Ho S, Kunz B, Hess K, Knott G et al. (2023) Dynamics of CLIMP-63 S-acylation control ER morphology. Nat Commun 14: 264

Sandoz PA, van der Goot FG (2015) How many lives does CLIMP-63 have? Biochem Soc Trans 43: 222–8

Schlaitz AL, Thompson J, Wong CC, Yates JR, 3rd, Heald R (2013) REEP3/4 ensure endoplasmic reticulum clearance from metaphase chromatin and proper nuclear envelope architecture. Dev Cell 26: 315–23

Shibata Y, Shemesh T, Prinz WA, Palazzo AF, Kozlov MM, Rapoport TA (2010) Mechanisms determining the morphology of the peripheral ER. Cell 143: 774–88

Shibata Y, Voeltz GK, Rapoport TA (2006) Rough sheets and smooth tubules. Cell 126: 435–9

Simovic-Lorenz M, Ernst A (2025) Chromothripsis in cancer. Nat Rev Cancer 25: 79–92

Slep KC, Vale RD (2007) Structural basis of microtubule plus end tracking by XMAP215, CLIP-170, and EB1. Mol Cell 27: 976–91

Smyth JT, Beg AM, Wu S, Putney JW, Jr., Rusan NM (2012) Phosphoregulation of STIM1 leads to exclusion of the endoplasmic reticulum from the mitotic spindle. Curr Biol 22: 1487–93

Turgay Y, Champion L, Balazs C, Held M, Toso A, Gerlich DW, Meraldi P, Kutay U (2014) SUN proteins facilitate the removal of membranes from chromatin during nuclear envelope breakdown. J Cell Biol 204: 1099–109

Uhlen M, Fagerberg L, Hallstrom BM, Lindskog C, Oksvold P, Mardinoglu A, Sivertsson A, Kampf C, Sjostedt E, Asplund A et al. (2015) Proteomics. Tissue-based map of the human proteome. Science 347: 1260419

Ungricht R, Klann M, Horvath P, Kutay U (2015) Diffusion and retention are major determinants of protein targeting to the inner nuclear membrane. J Cell Biol 209: 687–703

Vedrenne C, Klopfenstein DR, Hauri HP (2005) Phosphorylation controls CLIMP-63-mediated anchoring of the endoplasmic reticulum to microtubules. Mol Biol Cell 16: 1928–37

Voronina N, Wong JKL, Hubschmann D, Hlevnjak M, Uhrig S, Heilig CE, Horak P, Kreutzfeldt S, Mock A, Stenzinger A et al. (2020) The landscape of chromothripsis across adult cancer types. Nat Commun 11: 2320

Wang B, Zhao Z, Xiong M, Yan R, Xu K (2022) The endoplasmic reticulum adopts two distinct tubule forms. Proc Natl Acad Sci U S A 119: e2117559119

Wang Q, Crevenna AH, Kunze I, Mizuno N (2014) Structural basis for the extended CAP-Gly domains of p150(glued) binding to microtubules and the implication for tubulin dynamics. Proc Natl Acad Sci U S A 111: 11347–52

Wang T, Hanus C, Cui T, Helton T, Bourne J, Watson D, Harris KM, Ehlers MD (2012) Local zones of endoplasmic reticulum complexity confine cargo in neuronal dendrites. Cell 148: 309–21

Waterhouse AM, Procter JB, Martin DM, Clamp M, Barton GJ (2009) Jalview Version 2--a multiple sequence alignment editor and analysis workbench. Bioinformatics 25: 1189–91

Wei RR, Al-Bassam J, Harrison SC (2007) The Ndc80/HEC1 complex is a contact point for kinetochore-microtubule attachment. Nat Struct Mol Biol 14: 54–9

Weingarten MD, Lockwood AH, Hwo SY, Kirschner MW (1975) A protein factor essential for microtubule assembly. Proc Natl Acad Sci U S A 72: 1858–62

Westrate LM, Lee JE, Prinz WA, Voeltz GK (2015) Form follows function: the importance of endoplasmic reticulum shape. Annu Rev Biochem 84: 791–811

Wilkins BJ, Rall NA, Ostwal Y, Kruitwagen T, Hiragami-Hamada K, Winkler M, Barral Y, Fischle W, Neumann H (2014) A cascade of histone modifications induces chromatin condensation in mitosis. Science 343: 77–80

Wojcik E, Basto R, Serr M, Scaerou F, Karess R, Hays T (2001) Kinetochore dynein: its dynamics and role in the transport of the Rough deal checkpoint protein. Nat Cell Biol 3: 1001–7

Yang L, Guan T, Gerace L (1997) Integral membrane proteins of the nuclear envelope are dispersed throughout the endoplasmic reticulum during mitosis. J Cell Biol 137: 1199–210

Yu F, Courjaret R, Assaf L, Elmi A, Hammad A, Fisher M, Terasaki M, Machaca K (2024) Mitochondria-ER contact sites expand during mitosis. iScience 27: 109379

Zhao G, Liu S, Arun S, Renda F, Khodjakov A, Pellman D (2023) A tubule-sheet continuum model for the mechanism of nuclear envelope assembly. Dev Cell 58: 847–865 e10

Zheng P, Obara CJ, Szczesna E, Nixon-Abell J, Mahalingan KK, Roll-Mecak A, Lippincott-Schwartz J, Blackstone C (2022) ER proteins decipher the tubulin code to regulate organelle distribution. Nature 601: 132–138

